# Rapid and efficient *in vivo* astrocyte-to-neuron conversion with regional identity and connectivity?

**DOI:** 10.1101/2020.08.16.253195

**Authors:** Lei-Lei Wang, Carolina Serrano Garcia, Xiaoling Zhong, Shuaipeng Ma, Chun-Li Zhang

## Abstract

*In vivo* reprogramming of glia into functional neurons emerges as potential regeneration-based therapeutics for neural injuries or neurological diseases. Recent studies show that AAV-based manipulation of certain factors can rapidly and highly efficiently convert resident glia into functional neurons with brain region-specificity and precise connectivity. Using NEUROD1 as an example, we here show that the presumed astrocytes-converted neurons are essentially endogenous neurons in the adult mouse brain. AAV-mediated co-expression of NEUROD1 and a reporter indeed specifically, rapidly, and efficiently induces numerous reporter-labeled neurons. However, these neurons cannot be traced back to quiescent or reactive astrocytes by using stringent lineage-mapping strategies. Conversely, reporter-labeled neurons cannot be detected when NEUROD1 is strictly expressed in adult brain astrocytes. Through a retrograde labeling approach, our results rather reveal that endogenous neurons are the cell source for NEUROD1-induced reporter-labeled neurons. These results underline the indispensable value of stringent lineage-tracing strategies and beg for cautious interpretation of the *in vivo* reprogramming phenomena.

## INTRODUCTION

Neural injury or degeneration often leads to a loss of neurons and the disruption of functional neurocircuits. Regeneration of neurons could constitute an ideal therapeutic strategy since the regenerated neurons are immunocompatible to the host. Except for a few neurogenic niches (Zhao et al., 2008), however, the adult mammalian brain and spinal cord lack intrinsic capacity to produce new neurons under normal or pathological conditions.

With the advancements of cell fate reprogramming, *in vivo* conversion of resident glial cells to functional new neurons emerges as a promising strategy for neural regeneration (Barker et al., 2018; Chen et al., 2015; Li and Chen, 2016; Pereira et al., 2019; Smith and Zhang, 2015; Tai et al., 2020; Torper and Gotz, 2017; Wang and Zhang, 2018). This is largely accomplished by the ectopic expression of a single or a combination of fate-determining factors or through knockdown of a single gene (Grande et al., 2013; Guo et al., 2014; Liu et al., 2015; Niu et al., 2013; Qian et al., 2020; Torper et al., 2015; Zhou et al., 2020). Resident glial cells are ideal cell sources for fate-conversions, since they become reactive and can proliferate to replenish themselves in response to injury or neurodegeneration. While SOX2-mediated *in vivo* reprogramming passes through an expandable neural progenitor state that resembles the endogenous neurogenesis process (Niu et al., 2015; Niu et al., 2013; Su et al., 2014; Wang et al., 2016), the glia-to-neuron conversions by all other reprogramming strategies are direct without a proliferative intermediate (Brulet et al., 2017; Chen et al., 2020; Liu et al., 2020; Liu et al., 2015; Pereira et al., 2017; Torper et al., 2015; Wu et al., 2020; Zhou et al., 2020). Employing the AAV-mediated gene manipulations, recent studies further show that resident glial cells can be rapidly and very efficiently converted into mature neurons with brain region-specificity and precise connectivity (Chen et al., 2020; Liu et al., 2020; Liu et al., 2015; Mattugini et al., 2019; Pereira et al., 2017; Qian et al., 2020; Torper et al., 2015; Wu et al., 2020; Zhou et al., 2020). Most excitingly, the AAV-mediated direct reprogramming of glial cells shows therapeutic benefits under injury or degenerative conditions (Chen et al., 2020; Qian et al., 2020; Wu et al., 2020; Zhou et al., 2020). Such results, if confirmed, will revolutionize regenerative medicine.

During the past decade, we have conducted a series of *in vivo* screens for factors capable of reprogramming resident glia (Niu et al., 2013; Su et al., 2014; Wang et al., 2016). We notice that numerous endogenous neurons could have been readily misidentified as the glia-converted if stringent lineage-tracing methods were not employed. In this study, we systematically reexamined AAV-mediated direct astrocyte-to-neuron (AtN) conversion by using NEUROD1 as an example. Our results confirm that NEUROD1 can indeed rapidly and efficiently induce numerous reporter-labeled neurons in the adult mouse brain. However, these neurons do not come from resident astrocytes or reactive glial cells. Rather, they are endogenous neurons that are misidentified as the astrocyte-converted if based on the less stringent reporter system.

## METHODS

### Animals

Wildtype C57BL/6J mice and the following transgenic mouse lines were obtained from The Jackson Laboratory: *Aldh1l1-CreER*^*T2*^(stock #029655) (Srinivasan et al., 2016) and *R26R-YFP* (stock #006148) (Srinivas et al., 2001). Both male and female mice at 8 weeks and older were used for all experiments. All mice were housed under a controlled temperature and a 12-h light/dark cycle with free access to water and food in an animal facility at UT Southwestern. All animal procedures were approved by the Institutional Animal Care and Use Committee at UT Southwestern.

### AAV vectors and virus production

The following AAV vectors were constructed through PCR-based subcloning: *pAAV-hGFAP-GFP, pAAV-hGFAP-Cre, pAAV-hGFAP-mCherry, pAAV-hGFAP-NEUROD1-T2A-mCherry, pAAV-CAG-FLEX-mCherry*, or *pAAV-CAG-FLEX-NEUROD1-T2A-mCherry*. Unless indicated otherwise, the *hGFAP* promoter used in this study is the synthetic 681-bp *gfaABC1D* derived from the 2.2-kb *gfa2* promoter (Lee et al., 2008). Both promoters exhibit identical astrocyte-restricted expression patterns through transgenic analyses (Lee et al., 2008). The *AAV5-hGFAP*-Cre* virus was obtained from Addgene (#105550), in which Cre is driven by the 2.2-kb *gfa2* promoter. The *FLEX* vectors were based on *pAAV-FLEX-GFP* (Addgene #28304). *pAAV-CAG-GFP* was obtained from Addgene (#37825). All vectors were verified through DNA sequencing and restriction digestions. AAV virus was packaged with *pAd-deltaF6* (Addgene #112867) and the helper *pAAV2/5* (Addgene #104964), *pAAV2/8* (Addgene #112864), *pAAV2/9* (Addgene #112865), or *rAAV2-retro* (Addgene #81070) in HEK293T cells. Briefly, HEK293T cells were transfected with the packaging plasmids and a vector plasmid. Three days later, virus was collected from the cell lysates and culture media. Virus was purified through iodixanol gradient ultracentrifuge, washed with PBS and concentrated with 100K PES concentrator (Pierce™, Thermo Scientific). Viral titers were determined by quantitative PCR with ITR primers (forward: 5-*GGAACCCCTAGTGATGGAGTT*-3; reverse: 5-*CGGCCTCAGTGAGCGA*-3). Virus with a titer of 2e13 GC/mL was used for experiments.

### Brain injury and intracerebral injection

The controlled cortical impact (CCI) model of TBI was employed as previous described (Chen et al., 2019). Under anesthesia, a skin incision was made in the mouse forehead to expose the skull. A craniotomy was performed over the right hemisphere and the bone flap (2.2 mm in diameter) was carefully removed. A cortical injury (0 bregma, 1 mm lateral to the sagittal suture line) was introduced at an impact depth of 0.5 mm with a 2-mm diameter round impact tip (3.0 m/s speed and 100 ms dwell time) by using an electromagnetically driven CCI device (Impact One™ Stereotaxic CCI Instrument, Leica). After injury, skin was sutured and further secured with liquid adhesive. Intracerebral viral injections (1 µL per injection) were performed on a stereotaxic frame. The injection coordinates were as the following: +1.0 mm anterior/posterior (AP), ±1.5 mm medial/lateral (ML), and −0.8 mm dorsal/ventral from skull (DV) for uninjured cortex; 0 mm AP, ±1.5 mm ML, and −0.8 mm DV for injured cortex.

### Retrograde labeling of cortical motor neurons and intracerebral injection

*rAAV2-retro-CAG-GFP* virus (2 µL per mouse) was injected into the T6 level of dorsal spinal cord to target the corticospinal tract. Ten days post *rAAV2-retro* injection, we stereotaxically injected AAV5 virus into the cortex with the following coordinates: −0.5 mm AP, ±1.5 mm ML, and −1.0 mm DV.

### Tamoxifen and BrdU administration

Tamoxifen (10540-29-1; Cayman Chemical) was dissolved in a mixture of ethanol and sesame oil (1:9 by volume) at a concentration of 40 mg/mL. Tamoxifen was administered through intraperitoneal injections at a daily dose of 1 mg/10g body weight for 3-5 days. BrdU (H27260; Alfa Aesar Chemical; 0.5 g/L) was supplied in drinking water for durations as indicated in the text.

### Immunohistochemistry and quantification

Mice were euthanized and perfused with intracardial injection of 4% (w/v) paraformaldehyde in phosphate-buffered saline (PBS). Brains were isolated and post-fixed overnight with 4% (w/v) paraformaldehyde at 4°C. After cryoprotection with 30% sucrose in PBS for 72 h at 4 °C, 40-µm brain slices were collected through a sliding microtome (Leica). For immunostaining, brain sections were treated with 50% formamide in 1 X SSC buffer for 2 h at 65 °C. For BrdU staining, the sections were pretreated with 2N HCl for 30 min at 37 °C. The following primary antibodies were used: GFP (Chicken; 1:2000; Aves Labs), mCherry (Goat; 1:2000; Mybiosource), NeuN (Rabbit; 1:10000; Abcam), GFAP (Mouse; 1:2000; Sigma), Cre (Rabbit; 1:500; Covance), ALDH1L1 (Mouse; 1:200; Neuromab), ALDOC (Goat; 1:100; Santa Cruz), BrdU (Rat; 1:500; Bio-Rad). Alexa Fluor 488-, 555-, or 647-conjugated corresponding secondary antibodies from Jackson ImmunoResearch were used for indirect fluorescence (1:2000). Nuclei were counterstained with Hoechst 33342 (Hst). Images were captured using a Zeiss LSM700 confocal microscope. Four representative images from the top and middle layer of the cortex near the injection site were collected and quantified for each brain sample.

### Quantification of cell density

NeuN^+^ or GFAP^+^ cells were quantified from a 1-µm thick confocal image with an area of 0.102977 mm^2^. Four to six random confocal images surrounding the viral injection area were analyzed for each animal.

### Statistical analysis

Quantification data are presented as mean ± SEM from 3-5 mice per group. Statistical analysis was performed by homoscedastic two-tailed Student’s t-test using the GraphPad Prism software v. 6.0. A p value < 0.05 was considered significant. Significant differences are indicated by *p < 0.05, **p < 0.01, ***p < 0.001, and ****p < 0.0001.

## Acknowledgements

We thank members of the Zhang laboratory for reagents, Dr. Woo-ping Ge for discussion and comments. C.L.Z. is a W. W. Caruth, Jr. Scholar in Biomedical Research and supported by the Welch Foundation (I-1724), the Decherd Foundation, Kent Waldrep Foundation Center for Basic Research on Nerve Growth and Regeneration, and NIH Grants (NS099073, NS092616, NS111776, NS117065 and NS088095).

## RESULTS

### Rapid and highly efficient *in vivo* astrocyte-to-neuron conversion?

To target brain astrocytes through an AAV virus-based approach, we first compared the cell type-specificity of several commonly used AAV serotypes: AAV5, AAV8, and AAV9 (Brulet et al., 2017; Chen et al., 2020; Liu et al., 2020; Liu et al., 2015; Pereira et al., 2017; Torper et al., 2015). To further restrict gene expression in astrocytes, we used the synthetic 681-bp *gfaABC1D* human *GFAP* promoter (*hGFAP*) to drive gene expression (Lee et al., 2008). This promoter exhibits astrocyte-restricted expression as the 2.2-kb *gfa2* promoter in transgenic analysis (Lee et al., 2008). Cell type-specificity was examined by the GFP reporter after stereotaxic intracerebral injections of these AAVs. When examined at 4 days post virus injections (dpv), GFP was predominantly detected in GFAP^+^ cortical astrocytes for all these AAVs (Fig. 1A, C; 97.6 ± 1.2% for AAV8, 94.1 ± 0.9% for AAV9, and 92.1 ± 1.4% for AAV5). GFP expression in NeuN^+^ cortical neurons was minimal (2 ± 1.2%, 5.9 ± 0.9% and 7.9 ± 1.4% for AAV8, AAV9 and AAV5, respectively). However, cell type-specificity of these AAVs was dramatically different when examined at 14 dpv (Fig. 1B, C). Only about 62.4 ± 8.0% and 69.2 ± 8.0% of the virus-transduced cells were GFAP^+^ astrocytes for AAV8 and AAV9, respectively, whereas the neuronal fractions were correspondingly increased. Furthermore, both AAV8 and AAV9 showed spatial preferences with predominant neurons or astrocytes in different cortical regions (Fig. 1B). On the other hand, the cell type-specificity of AAV5 remained rather stable, with 90.4 ± 8.0% of reporter in astrocytes and 9.6 ± 8.0% in neurons (Fig. 1B and C). As such, AAV5 was selected for all subsequent experiments.

**Figure 1.**
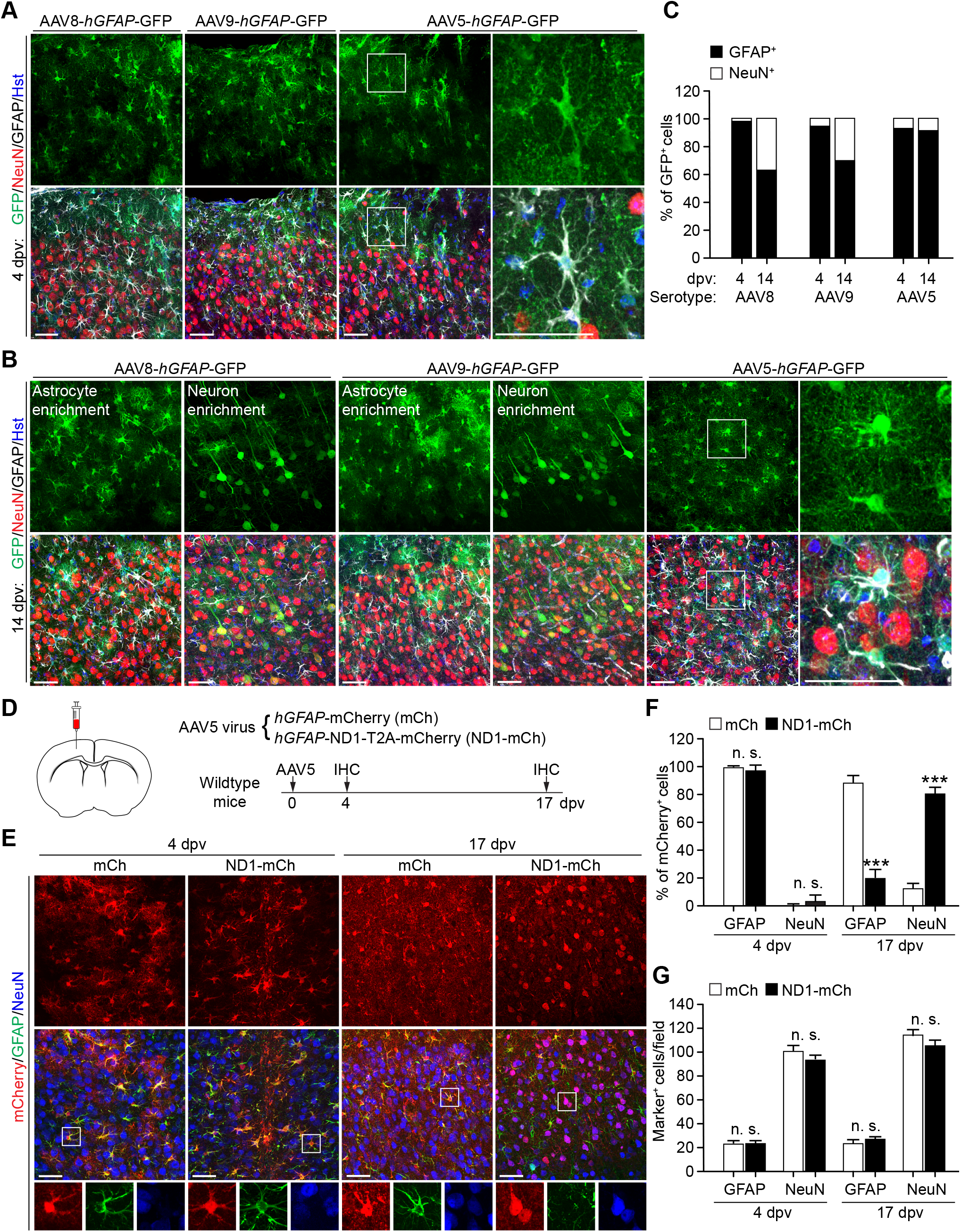
NEUROD1 induces rapid, specific, and highly efficient *in vivo* “AtN” conversion. (A) Confocal images showing marker expression in the virus-injected cortex. Samples were examined at 4 days post virus-injection (dpv). Enlarged views of the boxed regions are shown on the right. Scales, 50 µm. (B) Confocal images showing marker expression in the cortex at 14 dpv. Enlarged views of the boxed regions are shown on the right. Scales, 50 µm. (C) Cell type-specificity of the indicated AAV serotypes with AAV5 predominantly targeting astrocytes (n = 3-4 mice per serotype per time-point). (D) Experimental designs to examine “AtN” conversion by NEUROD1. IHC, immunohistochemistry; dpv, days post virus-injection. (E) Confocal images of the indicated markers at two time-points. Enlarged views of the boxed regions are shown at the bottom. Scales, 50 µm. (F) A time-course analysis showing time-dependent and highly efficient “AtN” conversion by NEUROD1 (n = 3-4 mice per group per time-point; ***p < 0.001 when compared to the *mCh* control; n.s., not significant). (G) Relatively stable cell densities surrounding the virus-injected cortical regions (n = 3-4 mice per group per time-point; n.s., not significant).

We then made AAV5 virus to express *mCherry* (*mCh*) or *NEUROD1-T2A-mCherry* (*ND1-mCh*) under the *hGFAP* promoter and stereotaxically injected into the cortex of adult wildtype mice (Fig. 1D). When examined at 4 dpv, the reporter mCherry was highly restricted to GFAP^+^ astrocytes for both viruses (Fig. 1E, F; 99.9 ± 0.1% for *mCh* and 96.4 ± 3.6% for *ND1-mCh*). The number of mCherry^+^ neurons was really small for these viruses at 4 dpv. Remarkably, when examined at 17 dpv, about 79.8 ± 4.4% of mCherry^+^ cells were NeuN^+^ neurons in the cortex of mice injected with the *ND1-mCh* virus (Fig. 1E, F). This was in sharp contrast to the control *mCh* virus-injected mice, in which 87.7 ± 5.0% of mCherry^+^ cells were still astrocytes. Such a result was largely consistent with what was previously reported on NEUROD1-mediated AtN conversion in adult mouse brains (Chen et al., 2020; Liu et al., 2020). Despite such rapid and robust appearance of mCherry^+^ neurons in the *ND1-mCh* group, however, we failed to detect significant changes on the density of either neurons or astrocytes in the virus-injected cortex (Fig. 1G). This was really unexpected as rapid and highly efficient AtN conversion would have resulted in an increased neuronal density and a correspondingly decreased astrocyte density.

### Reactive astrocytes are not a cell origin for NEUROD1-induced mCherry^+^ neurons

A therapeutic promise for *in vivo* cell fate reprogramming is to convert reactive astrocytes into neurons following traumatic neural injuries or under degenerative conditions (Li and Chen, 2016; Pereira et al., 2019; Smith et al., 2017; Smith et al., 2016; Tai et al., 2020; Wang and Zhang, 2018). We therefore examined whether those NEUROD1-induced mCherry^+^ neurons originated from reactive cortical astrocytes. Adult wildtype mice were subjected to traumatic brain injuries through controlled cortical impact (CCI) (Chen et al., 2019). Immediately following CCI, these mice were continuously administered with BrdU in drinking water to label proliferating cells including those reactive astrocytes. Seven days post injury, we injected either *mCh* or *ND1-mCh* AAV5 virus into the penumbra of the injured cortex and performed analyses after another 17 days (Fig. 2A). Approximately 71.5 - 75.7% of GFAP^+^ cells were also BrdU^+^ surrounding the injured cortex in both virus groups, indicating robust labeling of reactive astrocytes (Fig. 2B, C). Under this injury condition, 74.0 ± 3.9% of mCherry^+^ cells in the *ND1-mCh* group were NeuN^+^, comparing to 2.3 ± 1.7% in the control *mCh* group (Fig. 2D, E). However, only 2.0 ± 1.5% of mCherry^+^ cells were BrdU^+^NeuN^+^ in the *ND1-mCh* group (Fig. 2E), indicating a really low BrdU-labeling efficiency. Of note, many of these BrdU^+^NeuN^+^ cells exhibited a glial cell-like morphology with small nuclei, implicating induction of NeuN expression in glial cells (Fig. 2E). Once again, despite such a high percentage of mCherry^+^ cells being NeuN^+^ in the *ND1-mCh* group, quantification of neuronal density failed to detect a significant increase surrounding the injury penumbra (Fig. 2F). Together, these results indicate that reactive astrocytes are not a major cell source for NEUROD1-induced mCherry^+^ neurons.

**Figure 2.**
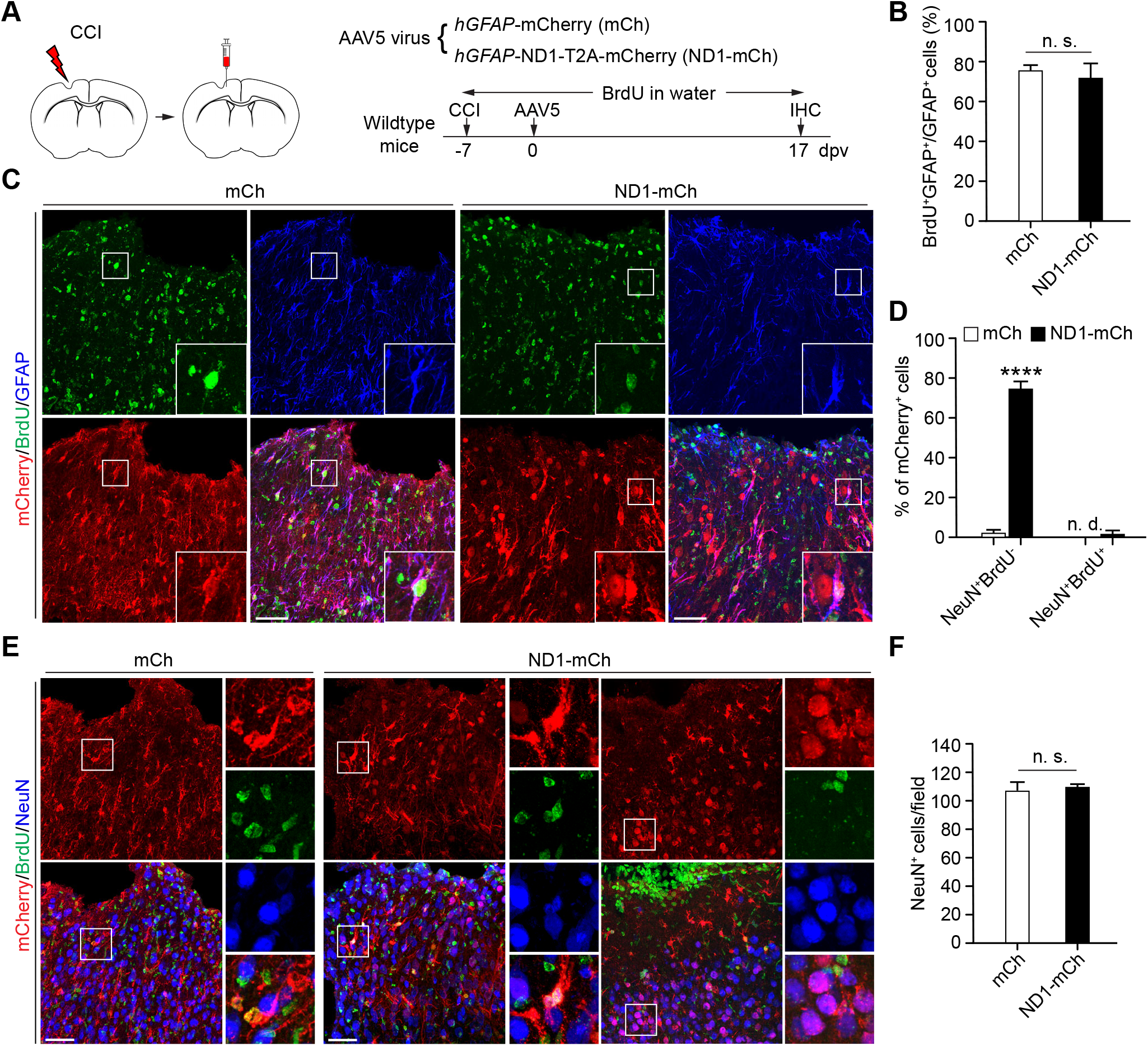
Reactive astrocytes are not involved in “AtN” conversion by NEUROD1. (A) Experimental designs to trace reactive glial cells. CCI, controlled cortical impact; IHC, immunohistochemistry; dpv, days post virus-injection. (B) High labeling-efficiency of reactive astrocytes by BrdU in drinking water (n = 4 mice per group; n.s., not significant). (C) Confocal images of BrdU-labeled astrocytes at 17 dpv. Enlarged views of the boxed regions are also shown. Scales, 50 µm. (D) Quantification showing barely any involvement of BrdU^+^ cells in “AtN” conversion by NEUROD1 at 17 dpv (n = 4 mice per group, ****p < 0.0001 when compared to the *mCh* control; n.d., not detected). (E) Confocal images showing the lack of BrdU-incorporation in neurons. A BrdU-labeled astrocyte-like NeuN^+^ cell is shown in the middle panel of *ND1-mCh* group. Scales, 50 µm. (F) Relatively stable neuronal densities surrounding the virus-injected cortical regions (n = 4 mice per group; n.s., not significant).

### The AAV-based Cre-FLEX system fails to restrict gene expression to astrocytes

The failure to label mCherry^+^ neurons with BrdU in the *ND1-mCh* group suggests that they might originate from quiescent astrocytes. To broadly target astrocytes in wildtype mice, we employed the AAV-based Cre-FLEX system by using two sets of AAV vectors (Fig. 3A), as previously described (Chen et al., 2020; Pereira et al., 2017; Torper et al., 2015). One AAV vector, *hGFAP-Cre*, expressed Cre recombinase under the *hGFAP* promoter to target astrocytes. The second set of AAV vectors, *F-mCh* and *F-ND1-mCh*, expressed mCherry and NEUROD1-T2A-mCherry in a Cre-dependent manner under the constitutively active CAG promoter.

**Figure 3.**
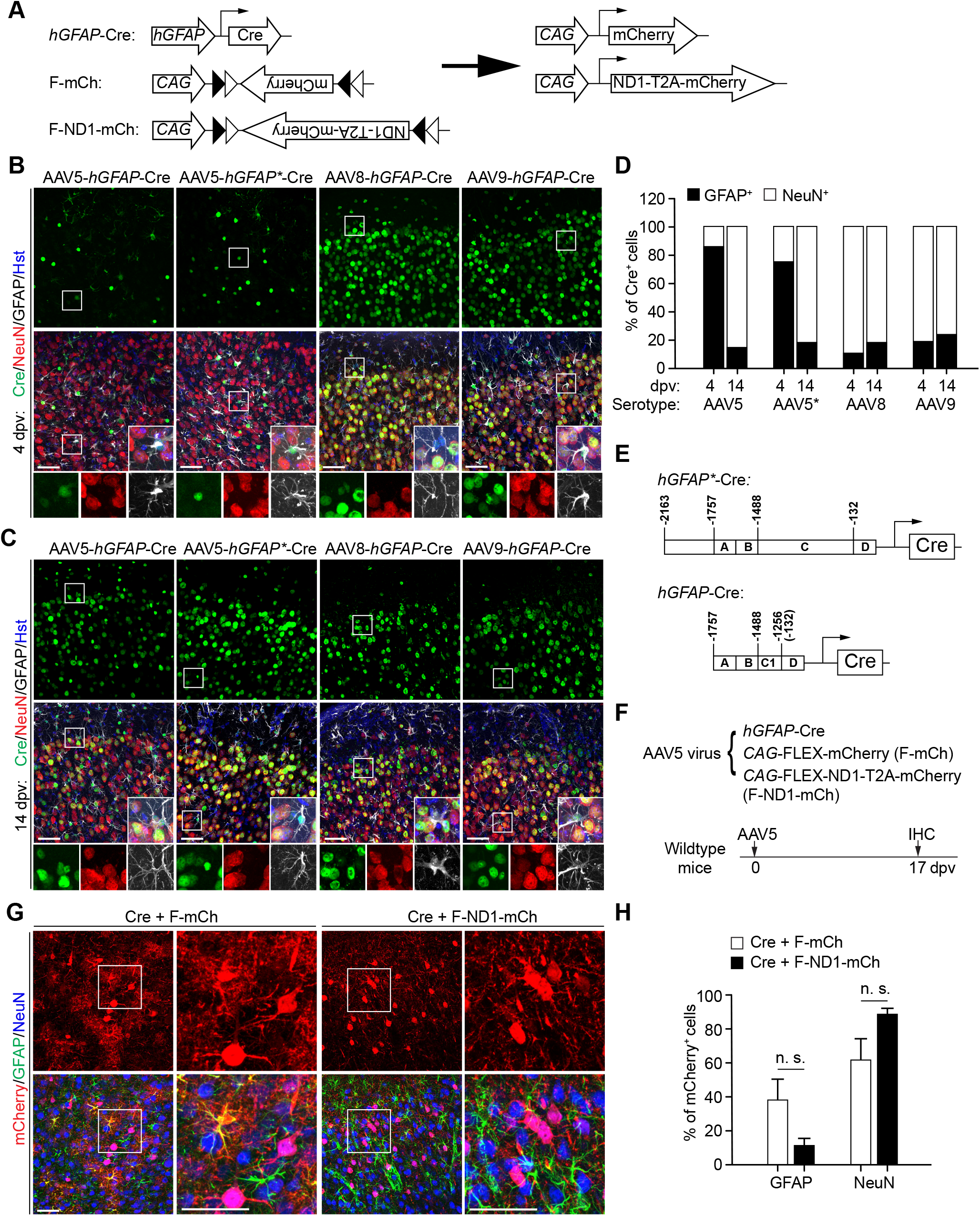
AAV-based Cre-FLEX system is unsuitable for examining “AtN” conversion. (A) A schematic illustration of the AAV-based Cre-FLEX system. (B) Cre expression in the virus-injected cortex at 4 dpv. Enlarged views of the boxed regions are also shown. Scales, 50 µm. (C) Cre expression in the virus-injected cortex at 14 dpv. Enlarged views of the boxed regions are also shown. Scales, 50 µm. (D) Quantification showing a lack of astrocyte-specificity for all the examined serotypes at a later time (n = 4 mice per serotype per time-point). (E) A schematic representation of the *GFAP* promoters used in this study. (F) Experimental designs to examine “AtN” conversion through the AAV-based Cre-FLEX system. (G) Confocal images of the indicated markers examined at 17 dpv. Enlarged views of the boxed regions are shown on the right panels for each group. Scales, 50 µm. (H) Quantification showing an unsuitability of the AAV-based Cre-FLEX system for examining “AtN” conversion (n = 4 mice per group; n.s., not significant).

We first examined cell type specificity of the *AAV-hGFAP-Cre* with the commonly used serotypes after stereotaxic injections. When examined at 4 and 14 dpv, both AAV8 and AAV9 serotypes showed Cre expression predominantly in NeuN^+^ neurons (Fig. 3B-D). The *AAV5-hGFAP-Cre* showed astrocyte-restricted expression at 4 dpv; however, neurons became the predominant cell type when examined at 14 dpv (Fig. 3B-D). We also examined the commercially available *AAV5*.*GFAP*.*Cre*.*WPRE*.*hGH* from Addgene (#105550; referred as *AAV5-hGFAP*-Cre* in this study), in which Cre is driven by the 2.2 kb *gfa2* human *GFAP* promoter (Lee et al., 2008) (Fig. 3E). Cell type analysis showed a similar pattern as the *AAV5-hGFAP-Cre* virus, with astrocytes and neurons as the predominant cells at 4 and 14 dpv, respectively (Fig. 3B-D). Despite these results, we co-injected *AAV5-hGFAP-Cre* virus with either *F-mCh* or *F-ND1-mCh* AAV5 virus into the cortical regions of adult wildtype mice. When examined at 17 dpv, mCherry^+^ neurons were observed in all the groups, though the *F-ND1-mCh* group showed less within-group variations (Fig. 3G-H; 61.8 ± 12.0% for *F-mCh* and 88.6 ± 3.5% for *F-ND1-mCh*). Together, these results indicate that the AAV-based Cre-FLEX system lacks the required cell type-specificity to trace *in vivo* AtN conversions.

### Astrocyte-restricted NEUROD1 expression fails to induce mCherry^+^ neurons

To specifically target endogenous astrocytes in adult mice, we employed the recently developed tamoxifen-inducible *Aldh1l1-CreER*^*T2*^ transgenic line (Srinivasan et al., 2016). We first confirmed cell type-specificity and labeling efficiency by crossing this line to the *R26R-YFP* reporter line (Fig. 4A). After 4 daily tamoxifen administrations, cortical regions were analyzed. The percentages of YFP^+^ cells expressing the astrocyte markers ALDH1L1, ALDOC, and GFAP were 95.7 ± 2.6%, 99.9 ± 0.1%, and 43.6 ± 6.4%, respectively (Fig. 4C). The lower percentage of GFAP-expressing YFP^+^ cells is expected, since GFAP is mainly expressed in reactive but not quiescent cortical astrocytes. Consistent with the previous report (Srinivasan et al., 2016), 4.3 ± 2.6% of YFP^+^ cells also expressed the neuronal marker NeuN; however, these cells had much lower level of YFP expression. The labeling efficiency was determined by the percentage of astrocytes being traced by YFP. They were 96.5 ± 1.8%, 94.1 ± 2.2%, and 94.3 ± 3.9% for ALDH1L1, ALDOC, and GFAP, respectively (Fig. 4D). These results confirm that the *Aldh1l1-CreER*^*T2*^ transgenic line can be employed to highly efficiently and specifically target endogenous astrocytes in adult mice (Srinivasan et al., 2016).

**Figure 4.**
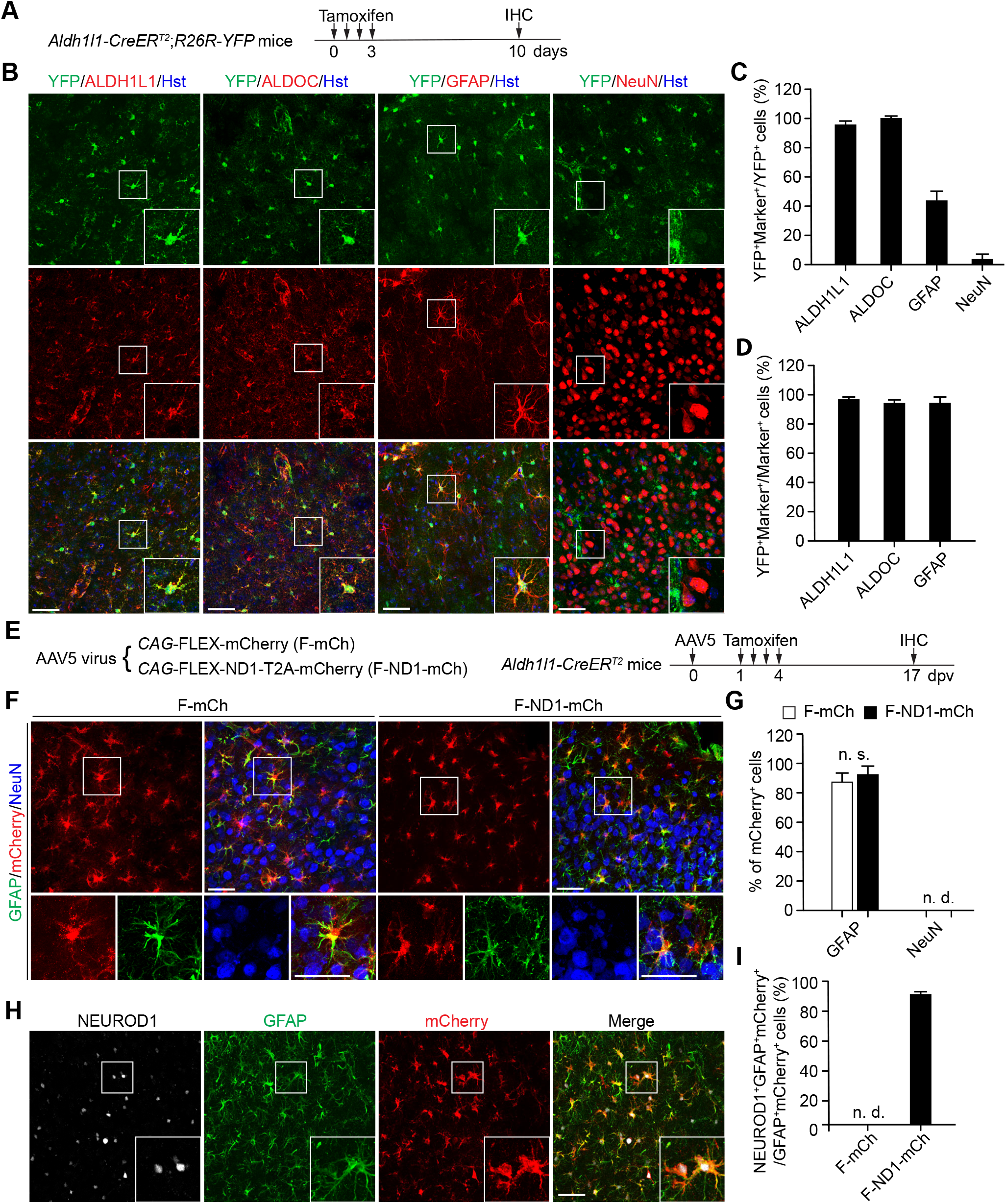
Astrocyte-restricted NEUROD1 fails to induce “AtN” conversion. (A) Experimental designs to examine tamoxifen-inducible Cre activity in adult astrocytes. (B) Confocal images of the tamoxifen-induced YFP reporter in the indicated cell types. Enlarged views of the boxed regions are also shown. Scales, 50 µm. (C) Quantification showing an astrocyte-restricted Cre activity (n = 4 mice). (D) Quantification showing a high labeling-efficiency of adult astrocytes by the tamoxifen-inducible approach (n = 4 mice). (E) Experimental designs to examine “AtN” conversion by astrocyte-restricted NEUROD1. (F) Confocal images showing a lack of the reporter mCherry in neurons at 17 dpv. Enlarged views of the boxed regions are also shown. Scales, 50 µm. (G) Quantification showing a failure of “AtN” conversion by astrocyte-restricted NEUROD1 at 17 dpv (n = 4 mice per group; n.s., not significant; n.d., not detected). (H) Confocal images showing robust NEUROD1 expression in astrocytes. Enlarged views of the boxed regions are also shown. Scales, 50 µm. (I) Quantification showing NEUROD1 expression in a majority of virus-transduced astrocytes at 17 dpv (n = 4 mice; n.d., not detected).

Adult *Aldh1l1-CreER*^*T2*^ mice were then intracerebrally injected with the Cre-dependent *F-mCh* or *F-ND1-mCh* AAV5 virus. These mice were treated with tamoxifen and analyzed at 17 dpv (Fig. 4E). Immunohistochemistry showed that the reporter mCherry was restricted to GFAP^+^ astrocytes in both experimental groups (Fig. 4F; 87.8 ± 6.4% for *F-mCh* and 92.8 ± 6.1% for *F-ND1-mCh*).

Unexpectedly, mCherry^+^ neurons were not detectable in the *F-ND1-mCh* group. To rule out the possibility that NEUROD1 might not be expressed in cortical astrocytes of mice injected with the *F-ND1-mCh* AAV5 virus, we performed immunostaining by using an antibody for NEUROD1. Confocal microscopy and quantification showed that 89.2 ± 2.4% of mCherry^+^ cells were stained positive for NEUROD1 (Fig. 4H, I). These results indicate that mCherry^+^ neurons cannot be induced by NEUROD1 when its expression is restricted to adult astrocytes through a tamoxifen-dependent transgenic approach.

### Neither tamoxifen nor genetic background affects NEUROD1’s ability to induce mCherry^+^ neurons

An alternative explanation for the above results was that tamoxifen might have abolished the reprogramming ability of NEUROD1 in adult *Aldh1l1-CreER*^*T2*^ mice. To examine such a possibility, we took a dual-reporter approach. One reporter was the Cre-dependent YFP in *Aldh1l1-CreER*^*T2*^;*R26R-YFP* mice, as shown in Figure 4A-D. The second reporter was the Cre-independent mCherry in either *mCh* or *ND1-mCh* AAV5 virus, as indicated in Figure 1D-G. Adult *Aldh1l1-CreER*^*T2*^;*R26R-YFP* mice were treated with tamoxifen, followed by intracerebral injections of either *mCh* or *ND1-mCh* AAV5 virus (Fig. 5A). When examined at 17 dpv, 73.3 ± 8.9% of mCherry^+^ cells were NeuN^+^ neurons in the *ND1-mCh* group (Fig. 5B, C), consistent with the results in wildtype mice (Fig. 1E, F). However, only 0.1 ± 0.1% of these mCherry^+^NeuN^+^ cells were also traced by YFP (Fig. 5D). We also examined a delayed tamoxifen-treatment scheme, in which tamoxifen was administered following the intracerebral injections of either *mCh* or *ND1-mCh* AAV5 virus into adult *Aldh1l1-CreER*^*T2*^;*R26R-YFP* mice (Fig. 5E). When examined at 28 dpv, 92.0 ± 1.1% of mCherry^+^ cells were mCherry^+^NeuN^+^ neurons in mice injected with the *ND1-mCh* AAV5 virus. Once again, only 1.8 ± 0.7% of these mCherry^+^NeuN^+^ were also labeled by YFP (Fig. 5F, G). Together, these results clearly indicate that neither tamoxifen-treatment nor the transgenic mouse background blocks the ability of NEUROD1 to induce mCherry^+^NeuN^+^ neurons. However, nearly all of the mCherry^+^NeuN^+^ cells cannot be genetically mapped to a cell source of resident astrocytes.

**Figure 5.**
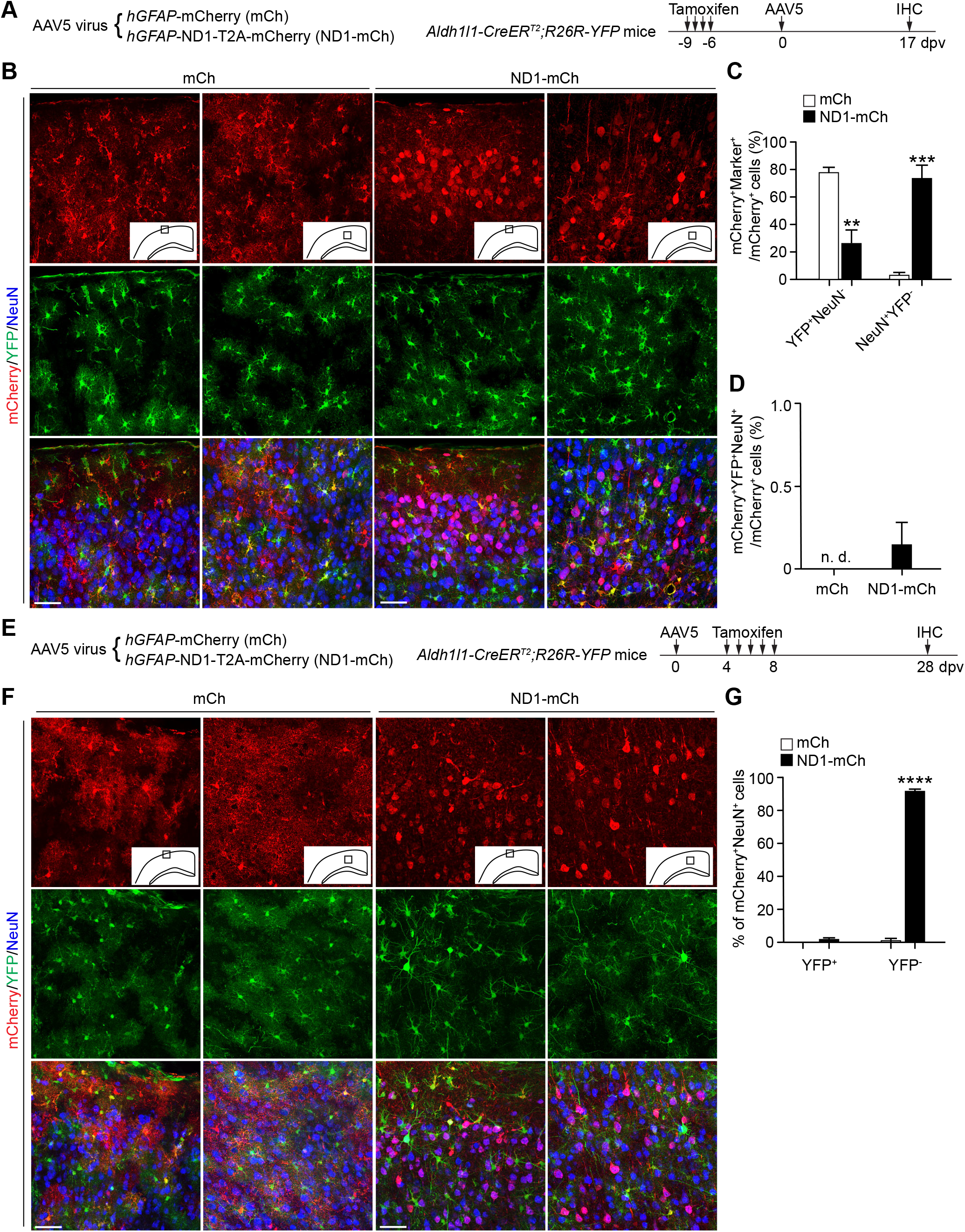
Neither tamoxifen nor genetic background affects “AtN” conversion by NEUROD1. (A) Experimental designs to examine the effect of tamoxifen on “AtN” conversion by NEUROD1 in lineage-tracing mice. Adult astrocytes are traced by the inducible YFP reporter. (B) Confocal images of the indicated two cortical regions for each group at 17 dpv. Scales, 50 µm. (C) Quantification showing robust “AtN” conversion but a lack of YFP-labeling of those neurons (n = 3-4 mice per group; **p = 0.0020 and ***p = 0.0003 when compared to the *mCh* control). (D) Quantification showing an extremely low YFP-labeling efficiency of NEUROD1-induced neurons (n = 4 mice per group). (E) Experimental designs to examine “AtN” conversion through an alternative tamoxifen-treatment scheme in lineage-tracing mice. Adult astrocytes are traced by the inducible YFP reporter. (F) Confocal images of the indicated two cortical regions for each group at 28 dpv. Scales, 50 µm. (G) Quantification showing robust “AtN” conversion but a lack of YFP-labeling of those neurons (n = 3 mice per group; ****p < 0.0001 when compared to the *mCh* control).

### Brain injury fails to precondition genetically traced astrocytes for conversion by NEUROD1

The environmental milieu may account for the failure of detecting AtN conversion by NEUROD1 in *Aldh1l1-CreER*^*T2*^;*R26R-YFP* mice, since injury or neurodegeneration may precondition resident glial cells for fate reprogramming (Grande et al., 2013; Guo et al., 2014; Heinrich et al., 2014). We examined this possibility by using the CCI model of traumatic brain injury (Chen et al., 2019). Resident astrocytes were genetically labeled with YFP in adult *Aldh1l1-CreER*^*T2*^;*R26R-YFP* mice after tamoxifen treatment. These mice were then subjected to CCI, followed by intracerebral injections of either *mCh* or *ND1-mCh* AAV5 virus near the injury penumbra (Fig. 6A). mCherry^+^NeuN^+^ cells could be observed in both experimental groups, with significantly more in the *ND1-mCh* group (Fig. 6B, C; 18.5 ± 5.4% for *mCh* and 64.9 ± 7.2% for *ND1-mCh*, **p = 0.002). Nonetheless, very few of them were YFP^+^ (Fig. 6B, C; 1.5 ± 1.5% for *mCh* and 2.5 ± 1.7% for *ND1-mCh*; p = 0.649). These mCherry^+^NeuN^+^YFP^+^ cells might be a result of labeling background, since 4.3 ± 2.6% of endogenous neurons can be genetically traced in *Aldh1l1-CreER*^*T2*^ mice (Srinivasan et al., 2016) (Fig. 4B, C). Despite a higher number of mCherry^+^NeuN^+^ cells in the *ND1-mCh* group, our measurement of neuronal density failed to show a significant difference between groups surrounding the injury penumbra (Fig. 6D).

**Figure 6.**
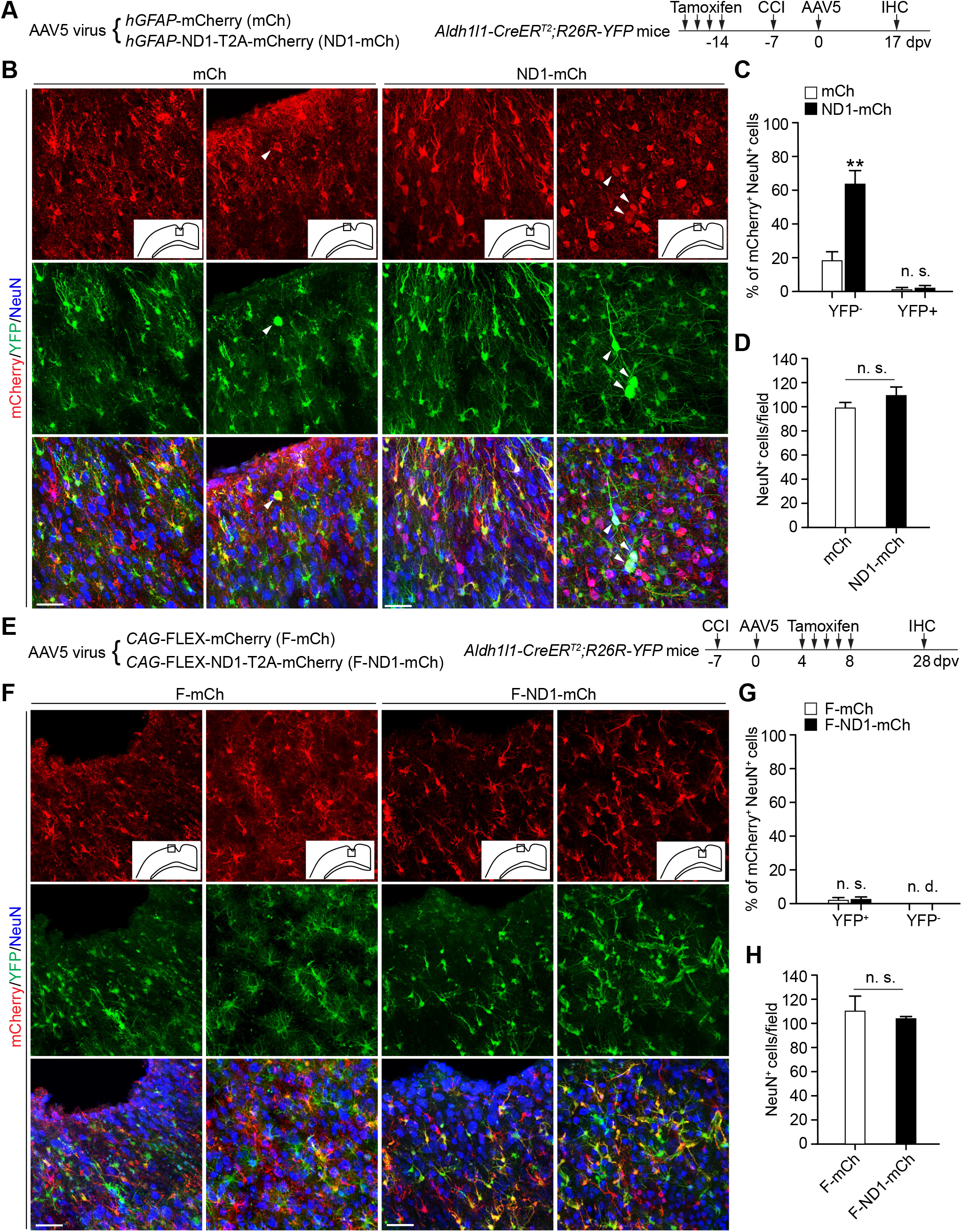
Brain injury fails to facilitate NEUROD1 to convert genetically traced astrocytes. (A) Experimental designs to examine the effect of brain injury on “AtN” conversion. Adult astrocytes are traced by the tamoxifen-induced YFP reporter. CCI, controlled cortical impact. (B) Confocal images of the indicated two cortical regions for each group at 17 dpv. A few YFP-labeled neurons, indicated by arrowheads, can be detected in each group after brain injury. Scales, 50 µm. (C) Quantification showing robust “AtN” conversion but a lack of YFP-labeling of those neurons (n = 4 mice per group; **p = 0.0022 when compared to the *mCh* control; n.s., not significant). (D) Quantification showing relatively stable neuronal densities surrounding the virus-injected cortical regions (n = 4 mice per group; n.s., not significant). (E) Experimental designs to examine “AtN” conversion by astrocyte-restricted NEUROD1. Adult astrocytes are traced by the inducible YFP reporter. (F) Confocal images of the indicated two cortical regions for each group at 28 dpv. Scales, 50 µm. (G) Quantification showing a lack of “AtN” conversion by astrocyte-restricted NEUROD1 at 28 dpv (n = 3-4 mice; n.s., not significant; n.d., not detected). (H) Quantification showing relatively stable neuronal densities surrounding the virus-injected cortical regions (n = 4 mice per group; n.s., not significant).

We also employed an experimental scheme by restricting the expression of NEUROD1 to genetically traced astrocytes after injury (Fig. 6E). Adult *Aldh1l1-CreER*^*T2*^;*R26R-YFP* mice were first subjected to CCI, followed by intracerebral injections of Cre-dependent *F-mCh* and *F-ND1-mCh* AAV5 virus. Astrocyte-restricted gene expression was initiated by tamoxifen treatment. When examined at 28 dpv, only 2.4 ± 1.2% and 2.9 ± 1.0% of mCherry^+^NeuN^+^ cells were also YFP^+^ for *F-mCh* and *F-ND1-mCh*, respectively (Fig. 6F, G; p = 0.749). Once again, such a lower number of triple positive cells might be a result of labeling background in *Aldh1l1-CreER*^*T2*^ mice (Srinivasan et al., 2016). On the other hand, mCherry^+^ but YFP^-^ neurons were not observed for either experimental group, excluding the possibility of selective silencing of the YFP reporter in the converted neurons (Fig. 6F, G). Consistently, no significant difference between groups was detected on neuronal density surrounding the injured cortex (Fig. 6H). Together, these results indicate that NEUROD1 is incapable of rapid and efficient AtN conversion even after traumatic brain injury.

### NEUROD1-induced mCherry^+^ neurons are endogenous neurons

Our results with stringent genetic lineage tracing and BrdU-labeling clearly indicate that resident astrocytes, reactive or quiescent, are not a cell source for NEUROD1-induced mCherry^+^ neurons in adult mice injected with the *ND1-mCh AAV5* virus. A key question was then what the cell source is for these NEUROD1-induced mCherry^+^ neurons. We finally explored the possibility that they might be endogenous neurons that were just misidentified as the astrocyte-converted. To examine such a possibility, we first labeled cortical motor neurons through the *rAAV2-retro*, an engineered AAV variant enabling retrograde access to projection neurons (Tervo et al., 2016). We injected the *rAAV2-retro-CAG-GFP* virus to target the cortical spinal tract (CST) at the 6^th^ thoracic spinal cord level (T6) of adult wildtype mice (Fig. 7A). When brains were examined at 10 and 28 dpv (Fig. 7B), a group of cortical motor neurons in the motor cortex were retrogradely labeled with GFP (Fig. 7C). Most importantly, GFP was not detected in any of the ALDH1L1^+^ or GFAP^+^ astrocytes in the motor cortex at both time points (Fig. 7D). Such a result showed extremely stringent cell type-specificity of the retrograde-labeling method. Adult wildtype mice were then first injected with the *rAAV2-retro-CAG-GFP* virus into the T6 CST, followed by intracerebral injections of *mCh* or *ND1-mCh* AAV5 virus into the motor cortex (Fig. 7E). When examined at 17 dpv, many mCherry^+^NeuN^+^ neurons were specifically detected in the *ND1-mCh* group (Fig. 7F, G). Of these, 37.3 ± 4.3% were also GFP^+^ surrounding the AAV5-injected cortex (Fig. 7F, H). This number was a huge underestimate of the contribution of endogenous neurons to the appearance of NEUROD1-induced mCherry^+^NeuN^+^ cells, since only a fraction of them were traced by the retrograde GFP reporter surrounding the AAV5-injected brain region. Together, these results show that endogenous neurons are the cell source for NEUROD1-induced mCherry^+^ neurons.

**Figure 7.**
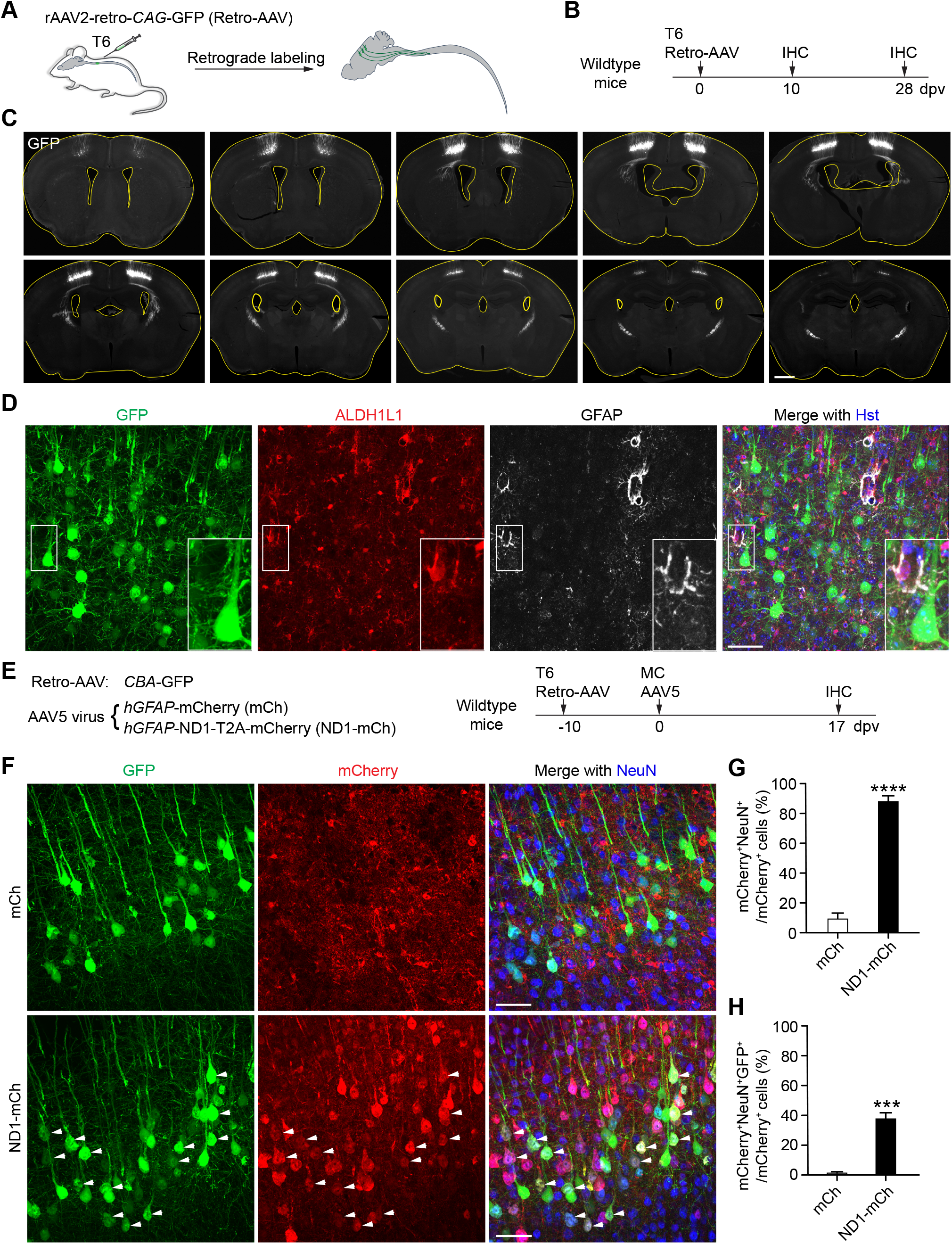
Endogenous neurons are the cell source for the presumed “AtN” conversion by NEUROD1. (A) A schematic on retrograde labeling of corticomotor neurons. T6, the 6^th^ thoracic spinal cord level. (B) Experimental designs to trace cortical motor neurons through retrograde labeling. (C) Series of brain sections showing specific labeling of the motor cortex by retro-AAV at 10 dpv. Ventricles are outlined. Scale, 1 mm. (D) Confocal images showing a complete lack of GFP-traced astrocytes in the motor cortex at 10 dpv. Scale, 50 µm. (E) Experimental designs to determine the contribution of endogenous neurons to NEUROD1-mediated “AtN” conversion. mc, motor cortex. (F) Confocal images of the motor cortex showing reporter-labeled neurons. Examples of neurons with dual reporters are indicated by arrowheads. Scales, 50 µm. (G) Quantification showing the presumed rapid and efficient “AtN” conversion by NEUROD1 in the cortex (n = 4-5 mice; ****p < 0.0001 when compared to the *mCh* control). (H) Quantification showing endogenous neurons as the cell source for the presumed “AtN” conversion (n = 4-5 mice; ***p = 0.0002 when compared to the *mCh* control).

## DISCUSSION

Using AAV-mediated co-expression of NEUROD1 and the mCherry reporter (Chen et al., 2020; Liu et al., 2020), our results confirm that NEUROD1 can specifically, rapidly, and highly efficiently induce numerous mCherry^+^ neurons in the adult mouse brain (Chen et al., 2020; Liu et al., 2020). Nonetheless, our multiple lineage-tracing strategies rather reveal that they are endogenous neurons but not converted from resident astrocytes.

First, continuous long-term BrdU-labeling after brain injury fails to show a contribution of reactive glial cells to the mCherry^+^ neurons. A potential caveat might be that BrdU is toxic to the newly converted neurons; however, this is not the case since new neurons from reprogrammed glia can be robustly traced by such a BrdU-labeling strategy (Niu et al., 2015; Niu et al., 2013; Su et al., 2014; Wang et al., 2016). Secondly, NEUROD1 fails to induce mCherry^+^ neurons when its expression is restricted to astrocytes of adult *Aldh1l1-CreER*^*T2*^ mice in a tamoxifen-inducible manner. One might argue that tamoxifen-treatments and the transgenic mouse background impede the reprogramming ability of NEUROD1; however, this is not the case since tamoxifen-independent NEUROD1 expression can induce numerous mCherry^+^ neurons in tamoxifen-treated *Aldh1l1-CreER*^*T2*^;*R26R-YFP* mice. Furthermore, these mCherry^+^ neurons cannot be traced by the astrocyte lineage reporter YFP, despite an approximate 95% labeling efficiency of astrocytes by YFP. Thirdly, NEUROD1 even fails to convert injury-preconditioned and genetically traced astrocytes. Fourthly, despite numerous mCherry^+^ neurons, the overall neuronal density remains unchanged in the NEUROD1 virus-injected regions, arguing against a net increase of neurons due to presumed AtN conversion. And finally, retrograde labeling clearly reveals that endogenous neurons are the cell source for NEUROD1-induced mCherry^+^ neurons. If any neurons were converted from astrocytes by AAV-mediated NEUROD1 expression, the number would be small, consistent with a previous report (Brulet et al., 2017). Our results, however, do not argue against a potential reprogramming capability of NEUROD1 under certain conditions, such as in cell culture or retrovirus-mediated expression after injury (Guo et al., 2014; Pataskar et al., 2016; Zhang et al., 2015).

An unresolved question is how NEUROD1 and potentially many other genetic manipulations, such as gene overexpression and knockdowns, specifically induce reporter expression in endogenous neurons. A possibility may be that these manipulations lead to a time-dependent activation of the *GFAP/Gfap* promoter in neurons. It has been observed that cell type-specificity of the *GFAP*/*Gfap* promoter can be altered by the expressed genes or under neurodegeneration (Hol et al., 2003; Su et al., 2004). Consistently, AAV5-mediated Cre expression switches from an initial astrocyte-restricted pattern to one predominantly in neurons, very different from that of either GFP or mCherry driven by the identical promoter. It awaits to be determined whether *Gfap* promoter activity is similarly influenced by the Cre-mediated manipulation of certain genes in the *GFAP-Cre line 77*.*6* transgenic mice (Gregorian et al., 2009). Another caveat working with constitutive Cre is that, when influenced by certain genetic manipulations, it might be transferred into neighboring neurons and induce reporter expression through exosomes or tunneling nanotubes (Fruhbeis et al., 2013; Wang et al., 2011). Future studies are clearly needed to understand how certain genetic manipulations, either overexpression or knockdowns, lead to altered cell type-specificity of gene promoters.

In conclusion, our results from using NEUROD1 as an example clearly indicate that stringent lineage-tracing strategies are indispensable when interpreting the results of *in vivo* neural reprogramming. Unlike cultured cells the fate of which can be directly observed, *in vivo* reprogramming occurs in a complex microenvironment often filled with numerous endogenous neurons. It is crucial to use well-controlled lineage mappings or time-lapse imaging to confirm the cell origin and/or exclude the possibility of misidentifying the endogenous neurons as the glia-converted.

## REFFERENCE

Barker, R.A., Gotz, M., and Parmar, M. (2018). New approaches for brain repair-from rescue to reprogramming. Nature 557, 329–334.

Brulet, R., Matsuda, T., Zhang, L., Miranda, C., Giacca, M., Kaspar, B.K., Nakashima, K., and Hsieh, J. (2017). NEUROD1 Instructs Neuronal Conversion in Non-Reactive Astrocytes. Stem cell reports 8, 1506–1515.

Chen, C., Zhong, X., Smith, D.K., Tai, W., Yang, J., Zou, Y., Wang, L.L., Sun, J., Qin, S., and Zhang, C.L. (2019). Astrocyte-Specific Deletion of Sox2 Promotes Functional Recovery After Traumatic Brain Injury. Cereb Cortex 29, 54–69.

Chen, G., Wernig, M., Berninger, B., Nakafuku, M., Parmar, M., and Zhang, C.L. (2015). In Vivo Reprogramming for Brain and Spinal Cord Repair. eNeuro 2.

Chen, Y.C., Ma, N.X., Pei, Z.F., Wu, Z., Do-Monte, F.H., Keefe, S., Yellin, E., Chen, M.S., Yin, J.C., Lee, G., et al. (2020). A NeuroD1 AAV-Based Gene Therapy for Functional Brain Repair after Ischemic Injury through In Vivo Astrocyte-to-Neuron Conversion. Mol Ther 28, 217–234.

Fruhbeis, C., Frohlich, D., Kuo, W.P., and Kramer-Albers, E.M. (2013). Extracellular vesicles as mediators of neuron-glia communication. Front Cell Neurosci 7, 182.

Grande, A., Sumiyoshi, K., Lopez-Juarez, A., Howard, J., Sakthivel, B., Aronow, B., Campbell, K., and Nakafuku, M. (2013). Environmental impact on direct neuronal reprogramming in vivo in the adult brain. Nat Commun 4, 2373.

Gregorian, C., Nakashima, J., Le Belle, J., Ohab, J., Kim, R., Liu, A., Smith, K.B., Groszer, M., Garcia, A.D., Sofroniew, M.V., et al. (2009). Pten deletion in adult neural stem/progenitor cells enhances constitutive neurogenesis. J Neurosci 29, 1874–1886.

Guo, Z., Zhang, L., Wu, Z., Chen, Y., Wang, F., and Chen, G. (2014). In vivo direct reprogramming of reactive glial cells into functional neurons after brain injury and in an Alzheimer’s disease model. Cell stem cell 14, 188–202.

Heinrich, C., Bergami, M., Gascon, S., Lepier, A., Vigano, F., Dimou, L., Sutor, B., Berninger, B., and Gotz, M. (2014). Sox2-mediated conversion of NG2 glia into induced neurons in the injured adult cerebral cortex. Stem cell reports 3, 1000–1014.

Hol, E.M., Roelofs, R.F., Moraal, E., Sonnemans, M.A., Sluijs, J.A., Proper, E.A., de Graan, P.N., Fischer, D.F., and van Leeuwen, F.W. (2003). Neuronal expression of GFAP in patients with Alzheimer pathology and identification of novel GFAP splice forms. Mol Psychiatry 8, 786–796.

Lee, Y., Messing, A., Su, M., and Brenner, M. (2008). GFAP promoter elements required for region-specific and astrocyte-specific expression. Glia 56, 481–493.

Li, H., and Chen, G. (2016). In Vivo Reprogramming for CNS Repair: Regenerating Neurons from Endogenous Glial Cells. Neuron 91, 728–738.

Liu, M.H., Li, W., Zheng, J.J., Xu, Y.G., He, Q., and Chen, G. (2020). Differential neuronal reprogramming induced by NeuroD1 from astrocytes in grey matter versus white matter. Neural Regen Res 15, 342–351.

Liu, Y., Miao, Q., Yuan, J., Han, S., Zhang, P., Li, S., Rao, Z., Zhao, W., Ye, Q., Geng, J., et al. (2015). Ascl1 Converts Dorsal Midbrain Astrocytes into Functional Neurons In Vivo. J Neurosci 35, 9336–9355.

Mattugini, N., Bocchi, R., Scheuss, V., Russo, G.L., Torper, O., Lao, C.L., and Gotz, M. (2019). Inducing Different Neuronal Subtypes from Astrocytes in the Injured Mouse Cerebral Cortex. Neuron 103, 1086–1095 e1085.

Niu, W., Zang, T., Smith, D.K., Vue, T.Y., Zou, Y., Bachoo, R., Johnson, J.E., and Zhang, C.L. (2015). SOX2 reprograms resident astrocytes into neural progenitors in the adult brain. Stem cell reports 4, 780–794.

Niu, W., Zang, T., Zou, Y., Fang, S., Smith, D.K., Bachoo, R., and Zhang, C.L. (2013). In vivo reprogramming of astrocytes to neuroblasts in the adult brain. Nature cell biology 15, 1164–1175.

Pataskar, A., Jung, J., Smialowski, P., Noack, F., Calegari, F., Straub, T., and Tiwari, V.K. (2016). NeuroD1 reprograms chromatin and transcription factor landscapes to induce the neuronal program. EMBO J 35, 24–45.

Pereira, M., Birtele, M., and Rylander Ottosson, D. (2019). Direct reprogramming into interneurons: potential for brain repair. Cell Mol Life Sci 76, 3953–3967.

Pereira, M., Birtele, M., Shrigley, S., Benitez, J.A., Hedlund, E., Parmar, M., and Ottosson, D.R. (2017). Direct Reprogramming of Resident NG2 Glia into Neurons with Properties of Fast-Spiking Parvalbumin-Containing Interneurons. Stem cell reports 9, 742–751.

Qian, H., Kang, X., Hu, J., Zhang, D., Liang, Z., Meng, F., Zhang, X., Xue, Y., Maimon, R., Dowdy, S.F., et al. (2020). Reversing a model of Parkinson’s disease with in situ converted nigral neurons. Nature 582, 550–556.

Smith, D.K., He, M., Zhang, C.L., and Zheng, J.C. (2017). The therapeutic potential of cell identity reprogramming for the treatment of aging-related neurodegenerative disorders. Prog Neurobiol 157, 212–229.

Smith, D.K., Wang, L., and Zhang, C.L. (2016). Physiological, pathological, and engineered cell identity reprogramming in the central nervous system. Wiley Interdiscip Rev Dev Biol 5, 499–517.

Smith, D.K., and Zhang, C.L. (2015). Regeneration through reprogramming adult cell identity in vivo. Am J Pathol 185, 2619–2628.

Srinivas, S., Watanabe, T., Lin, C.S., William, C.M., Tanabe, Y., Jessell, T.M., and Costantini, F. (2001). Cre reporter strains produced by targeted insertion of EYFP and ECFP into the ROSA26 locus. BMC Dev Biol 1, 4.

Srinivasan, R., Lu, T.Y., Chai, H., Xu, J., Huang, B.S., Golshani, P., Coppola, G., and Khakh, B.S. (2016). New Transgenic Mouse Lines for Selectively Targeting Astrocytes and Studying Calcium Signals in Astrocyte Processes In Situ and In Vivo. Neuron 92, 1181–1195.

Su, M., Hu, H., Lee, Y., d’Azzo, A., Messing, A., and Brenner, M. (2004). Expression specificity of GFAP transgenes. Neurochem Res 29, 2075–2093.

Su, Z., Niu, W., Liu, M.L., Zou, Y., and Zhang, C.L. (2014). In vivo conversion of astrocytes to neurons in the injured adult spinal cord. Nat Commun 5, 3338.

Tai, W., Xu, X.M., and Zhang, C.L. (2020). Regeneration Through in vivo Cell Fate Reprogramming for Neural Repair. Front Cell Neurosci 14, 107.

Tervo, D.G., Hwang, B.Y., Viswanathan, S., Gaj, T., Lavzin, M., Ritola, K.D., Lindo, S., Michael, S., Kuleshova, E., Ojala, D., et al. (2016). A Designer AAV Variant Permits Efficient Retrograde Access to Projection Neurons. Neuron 92, 372–382.

Torper, O., and Gotz, M. (2017). Brain repair from intrinsic cell sources: Turning reactive glia into neurons. Prog Brain Res 230, 69–97.

Torper, O., Ottosson, D.R., Pereira, M., Lau, S., Cardoso, T., Grealish, S., and Parmar, M. (2015). In Vivo Reprogramming of Striatal NG2 Glia into Functional Neurons that Integrate into Local Host Circuitry. Cell reports 12, 474–481.

Wang, L.L., Su, Z., Tai, W., Zou, Y., Xu, X.M., and Zhang, C.L. (2016). The p53 Pathway Controls SOX2-Mediated Reprogramming in the Adult Mouse Spinal Cord. Cell reports 17, 891–903.

Wang, L.L., and Zhang, C.L. (2018). Engineering new neurons: in vivo reprogramming in mammalian brain and spinal cord. Cell and tissue research 371, 201–212.

Wang, Y., Cui, J., Sun, X., and Zhang, Y. (2011). Tunneling-nanotube development in astrocytes depends on p53 activation. Cell Death Differ 18, 732–742.

Wu, Z., Parry, M., Hou, X.Y., Liu, M.H., Wang, H., Cain, R., Pei, Z.F., Chen, Y.C., Guo, Z.Y., Abhijeet, S., et al. (2020). Gene therapy conversion of striatal astrocytes into GABAergic neurons in mouse models of Huntington’s disease. Nat Commun 11, 1105.

Zhang, L., Yin, J.C., Yeh, H., Ma, N.X., Lee, G., Chen, X.A., Wang, Y., Lin, L., Chen, L., Jin, P., et al. (2015). Small Molecules Efficiently Reprogram Human Astroglial Cells into Functional Neurons. Cell stem cell 17, 735–747.

Zhao, C., Deng, W., and Gage, F.H. (2008). Mechanisms and functional implications of adult neurogenesis. Cell 132, 645–660.

Zhou, H., Su, J., Hu, X., Zhou, C., Li, H., Chen, Z., Xiao, Q., Wang, B., Wu, W., Sun, Y., et al. (2020). Glia-to-Neuron Conversion by CRISPR-CasRx Alleviates Symptoms of Neurological Disease in Mice. Cell 181, 590–603 e516.

